# A first description of the ichthyofauna of the Namibian Islands Marine Protected Area using baited remote underwater video systems

**DOI:** 10.64898/2026.09.14.751173

**Authors:** Ruth H. Leeney, Lauren de Vos, Mabuta A. Simataa, Angus van Wyk, Hayley Brand, Margit R. Wilhelm, Anthony Bernard

**Affiliations:** Sruth, Cluain Sceach, Dublin, Ireland; WILDTRUST, PO Box 21450, Mayors Walk, Pietermaritzburg 3208, South Africa; University of Namibia, Department of Fisheries and Ocean Science, Henties Bay, Namibia; Ministry of Agriculture, Fisheries, Water and Land Reform, Swakopmund, Namibia; South African Institute for Aquatic Biodiversity, Makhanda, South Africa; Department of Ichthyology and Fisheries Science, Rhodes University, Makhanda, South Africa; Starfish Street, Swakopmund, Namibia

**Keywords:** Benguela Current, biodiversity, BRUVs, elasmobranchs, *Chelidonichthys capensis*, *Galeorhinus galeus*, *Notorynchus cepedianus*, *Haploblepharus pictus*, monitoring

## Abstract

Data on the diversity, relative abundance and distribution of the ichthyofauna of Namibia’s only marine protected area, the Namibian Islands Marine Protected Area (NIMPA), are limited and there is no active monitoring programme in place. Generating a baseline for these species is essential to inform management plans and policy. Baited remote underwater video system (BRUVs) surveys were conducted between 2022 and 2024 in the central eastern NIMPA. Data from 114 BRUVs samples, collected in water less than 45 m deep, were analysed in this study. Of the 17 fish species recorded, five were elasmobranchs, and three fish species - *Galeorhinus galeus* (CR), *Notorynchus cepedianus* (VU) and *Argyrosomus inodorus* (VU) - are classified as threatened on the IUCN Red List. The most frequently recorded elasmobranch, *Haploblepharus pictus*, was ubiquitous and typified all habitat types and depth categories. The most frequently recorded actinopterygian fish (*Clinus spp.*) was sighted at 37% of sites. Depth was an important predictor of fish species abundance and community composition, but habitat was not. Both ichthyofaunal relative abundance and diversity appeared lower than in similar surveys in neighbouring South Africa and other temperate waters. Future research should aim to identify the drivers of the observed patterns. The findings provide an important baseline for long-term monitoring of the NIMPA, necessary to better describe biodiversity and inform the development of management strategies.

## 1. INTRODUCTION

Marine Protected Areas (MPAs) have been designated worldwide to protect local marine resources, mostly as a tool for fisheries management, habitat protection, and conservation of biodiversity (Guidetti et al. 2014; Sciberras et al. 2015; Kriegl et al. 2021). Whilst MPAs have been documented to contribute to the protection and recovery of teleost populations (e.g. Ojeda-Martinez et al. 2007; Rojo et al. 2019), their contribution to the protection of elasmobranchs is not as well understood. Some evidence of MPAs benefitting elasmobranchs exists (e.g. Albano et al. 2021; Goetze et al. 2024), but the outcomes can be variable and the extent of the protection may not be sufficient, especially when MPAs are isolated or of insufficient size (Dwyer et al. 2020; Faure-Beaulieu et al. 2024).

The Benguela Current Large Marine Ecosystem (BCLME) is one of the world’s major productive eastern-boundary currents, and the intense coastal upwellings in this system support plankton production, which in turn supports rich pelagic and demersal fish populations (Sumaila & Vasconcellos 2000; Boyer and Hampton 2001). Namibian fisheries are relatively low in diversity, but the BCLME supported what was one of the most productive marine ecosystems globally (Cochrane et al. 2009). Upwelling zones along the coast supported rich pelagic resources of Southern African sardine *(Sardinops sagax*), South African anchovy (*Engraulis capensis*), Cape horse mackerel (*Trachurus capensis*) and other pelagic fishes, whilst amongst the demersal species, Cape hakes (*Merluccius capensis* and *M. paradoxus*) were the most valuable and abundant (Sumaila 2000; Sumaila & Vasconcellos 2000; Willemse & Pauly 2004; Wilhelm et al. 2015). However, Namibian waters were heavily exploited, both legally, during the colonial era, and illegally thereafter. Heavy and unsustainable exploitation by ‘foreign’ fleets (including those of South Africa whilst Namibia was a South African colony) in the 1960s and 1970s resulted in the collapse of sardine and Cape hake stocks, two of the most important stocks in the country (Crawford et al. 1987; Belhabib et al. 2015). This has resulted in a degraded marine ecosystem characterised by decreased productivity, and likely had implications throughout the food web, for meso- and top-predators (Roux et al. 2013; Erasmus et al. 2021). Despite post-independence (from 1990 onwards) efforts for national control of fisheries in Namibian waters, most remaining fish stocks are considered either fully- or over-exploited (Belhabib et al. 2019). The fishing industry currently consists largely of industrial fleets, primarily bottom trawlers, mid-water trawlers and longliners, while the small-scale fishing sector is informal and small in size, consisting mainly of shore-based angling and seine netting (Sowman & Cardoso 2010). A licensing condition has prohibited trawling inside the 200-metre isobath since 1990, effectively protecting inshore waters from trawling (Wilhelm et al. 2015). Recreational fishing, primarily from the coast but also (to a lesser extent) from boats, is a popular pastime in Namibia and is the only inshore fishing activity occurring along certain parts of the coast. Some of the main species targeted by this recreational fishery – silver kob *Argyrosomus inodorus* and West coast steenbras *Lithognathus aureti* – are considered overfished (Gusha et al. 2025).

In addition to teleosts, baseline information on the diversity, distribution, and abundance estimates of elasmobranchs is crucial for the development of effective management and conservation initiatives (Garla et al. 2006; White et al. 2013). However, collecting data on these species can be challenging because most are highly mobile and many have broad geographic ranges (Dulvy et al. 2008; McCauley et al. 2012). Baseline data on elasmobranchs are especially lacking in many low-income countries, where resources for research and monitoring of marine species may be prioritised for those of high commercial value (RH Leeney pers. obs.). However, these species are critical components of healthy marine ecosystems and in some low-income countries, where they are targeted as sources of food and fins, they can be particularly important to the socio-economic wellbeing of small-scale fishing communities (e.g. Seidu et al. 2022; Leeney et al. 2018). Until recently, there was little research focus on non-commercial fish species in Namibian waters and as a result, the diversity, distribution and conservation status of these species is poorly understood. Generating a baseline for key fish species is essential to inform management and to design longer-term monitoring programmes that can detect changes in the status of species of conservation concern.

Surveys using baited remote underwater video systems (hereafter BRUVs) are increasingly being used to sample the relative abundance of fish assemblages in marine ecosystems (McLaren et al. 2015; Whitmarsh et al. 2017). Using these systems allows for the population to be sampled in a non-extractive way and allows for simultaneous counts of multiple taxa. BRUVs are especially useful for sampling in a wide range of depths and they have little impact on the area being studied, making them ideal for use in MPAs (Langlois et al. 2018). Results from studies comparing underwater visual census (UVC) with BRUVs have varied considerably, with recorded species richness sometimes being higher using BRUVs (e.g. La Manna et al. 2021) and in other cases, higher using UVC (e.g. Colton & Swearer 2010). BRUVs often record more cryptic species and mobile or transitory predatory species that are less likely to be detected during UVC (Colton & Swearer 2010; Davis et al. 2019; Cheal et al. 2021; Rolim et al. 2022). The use of BRUVs also lowers the risk of incorrect fish identifications and inter-observer variability by recording a permanent and reviewable record (Langlois et al. 2018).

A good understanding of the marine habitats and biodiversity in the Namibian Islands Marine Protected Area (NIMPA) is lacking. Such information is necessary to inform the management of the NIMPA, and ongoing monitoring is essential as a means of assessing whether the MPA is providing any protection to key species and habitats. The objectives of this study were to pilot the use of BRUVs in Namibian waters for the first time; to document the marine fauna in the NIMPA and to assess the relative abundance of species at sites throughout the NIMPA. The resulting dataset provides valuable insight into habitat and seabed types and marine faunal assemblages at the sampled sites. Piloting the use of BRUVs also allowed for an assessment of their value for monitoring marine fauna in Namibian waters.

## 2. MATERIALS & METHODS

### 2.1 Study area

The NIMPA is Namibia’s only MPA and is the second largest marine reserve off the African coast, encompassing c. 9,400 km^2^ of ocean (Fig. 1). Proclaimed in 2008 (Government Gazette of the Republic of Namibia 2009), it was designated primarily because of its importance for numerous seabird species; 11 species that are endemic to southern Africa breed on the islands within the MPA boundary (Kemper et al. 2007; Ludynia et al. 2012). It is located within the Lüderitz Upwelling Cell in the central region of the Benguela Current Large Marine Ecosystem (BCLME), where high levels of productivity historically supported high abundances of fish and predator species (Cochrane et al. 2009). Between its establishment in 2009 and the present, no formal management plan has been implemented for the NIMPA, although at the time of writing (February 2026), a draft management plan is in review with the Namibian Government.

**Figure 1:**
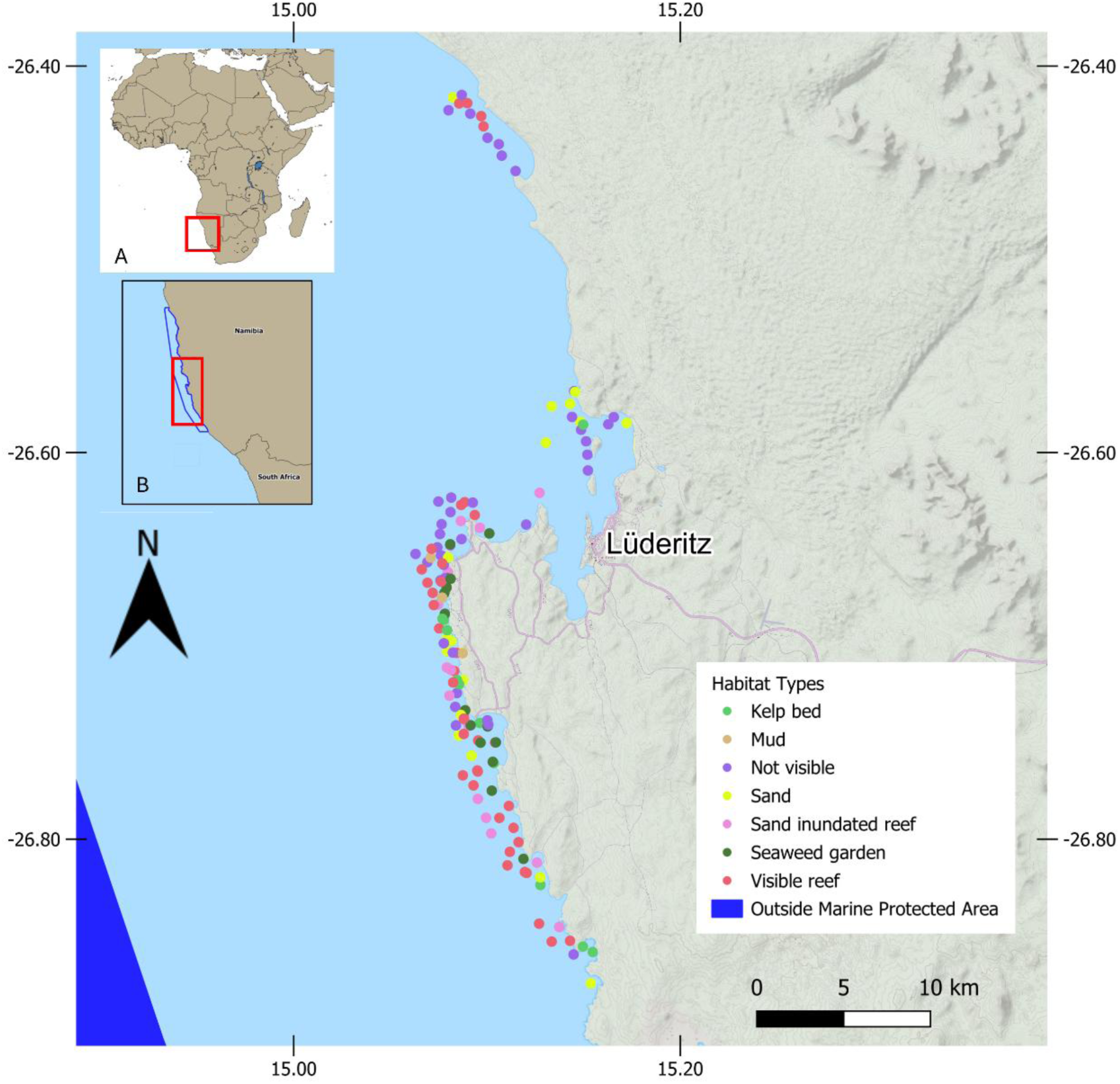
The distribution of 143 BRUVs samples collected during the study, colour-coded by habitat type. The area of light blue shading falls inside the Namibian Island Marine Protected Area (NIMPA) boundaries; the area of dark blue shading indicates waters outside the MPA boundary. The inset maps show (A) the location of the Namibian coast on the African continent, and (B) the survey area denoted in a red box, in the context of the location and boundaries of the NIMPA on Namibia’s coastline, delineated by the blue line.

The NIMPA encompasses a variety of marine and coastal habitats along Namibia’s coastline, including lagoons, wetlands, salt pans, rocky shores, reefs, sandy beaches, kelp forests and small islands (MARISMA EBSA Workstream 2023). The islands within the NIMPA have been modified through anthropogenic impacts such as intensive guano scraping activities (Griffiths et al. 2005) which, alongside the depletion of small pelagic fish stocks, has impacted some seabird populations (e.g. Pichegru et al. 2024). Historically, the biodiversity in the inter-tidal zones of the islands tended to be greater than elsewhere in the area, possibly due to significant nutrient input from seabird guano (MARISMA EBSA Workstream 2023) and thus guano scraping may also have impacted the intertidal and sub-tidal communities around the islands. Industrial fishing is not permitted inside the NIMPA, and the only fisheries currently permitted are recreational angling, recreational collection of West coast rock lobster (*Jasus lalandii*) and several small-scale fisheries using pots or traps and targeting crab, rock lobster and octopus, which likely have low levels of threatened species bycatch. The Namibian fishery for snoek (*Thyrsites atun*), based in Walvis Bay, fishes in surface waters and uses handlines with lures. In the past, blue shark (*Prionace glauca*) was occasionally caught but this appears to be uncommon now (N. Dreyer, pers. comm., in Leeney 2024a). Snoek vessels are not permitted to fish inside the NIMPA but anecdotal evidence suggests that some do. Prior to 2022, limited research had been conducted to describe the ichthyofaunal diversity within the NIMPA, except for the monitoring of species of commercial interest.

### 2.2 Study design

The lack of baseline data on bathymetry and seabed types in the NIMPA prevented any *a priori* development of a stratified sampling design to sample equally at different depth ranges and in different habitat types. A haphazard sampling approach (a non-statistical, non-probability technique where, in this case, sites were selected based on convenience or judgment without following a structured procedure) was therefore used, with the aim of collecting samples from a range of depths and habitat types. Once a general area was selected for sampling, the water depth was determined using the boat’s echosounder, which allowed the team to deploy BRUVs at a range of depths each day. Bottom type was only distinguishable on the echosounder as either reef or soft sediment and this limited information, as well as the presence of kelp at the surface, was used to attempt to distribute samples amongst different habitat types.

### 2.3 Sampling technique

Sampling was conducted from a 5 m, fibreglass boat suitable for the deployment and retrieval of the BRUVs. The methods used followed protocols described in detail in Langlois et al. (2020) and used extensively in South Africa (e.g. De Vos et al. 2014; Osgood et al. 2019; Martinez et al. 2024). Samples were collected only during daylight hours, typically between 08:00 and 16:00 Namibian time. One kg of crushed sardines (*Sardinops sagax*) was used as bait. Each BRUVs bore two GoPro 8 cameras (stereo systems were used; only the left camera’s footage was analysed for this study) and a central underwater housing containing a white LED light. After deployment, each system was left on the seabed to record for 60 minutes, after which it was retrieved, the camera batteries and memory cards were replaced, the bait container was refilled and the system was redeployed at a new site. The light was on for all deployments, and each light’s batteries were replaced after every three deployments of each system. Concurrent BRUVs deployments were kept at least 400 m apart to avoid bait plume overlap (Bond et al. 2018; Langlois et al. 2020) and the risk of the same animals being detected on more than one system. On the vessel, the following data were recorded for each deployment: a unique code for each sample, deployment location (GPS coordinates), deployment time, water depth and retrieval time.

Fieldwork was conducted in August 2022 (pilot field trip), February 2023 and February 2024. Two BRUVs samples were collected in August 2023, but poor sea conditions over a sustained period of time prevented the completion of the second winter field trip.

### 2.4 Video and data analysis

BRUVs footage was analysed in the SeaGIS EventMeasure software (seagis.com.au). The analysis period was 60 minutes from the time at which the BRUVs settled on the seafloor. All marine fauna were identified to the lowest possible taxon. Teleosts in the genus *Clinus* are considered highly cryptic and were identified as *Clinus* spp. For species that were not familiar to the analysts, a video clip in which the species appeared was made and was shared with biologists in Namibia and South Africa, and several recreational fishers in Namibia for identification. Each video was reviewed and analysed by at least two analysts to allow for all species identifications and counts (MaxN) of large groups of species to be verified.

For each deployment, the habitat type was classified based on a pre-defined list of habitats (Appendix I). Of the 143 sites sampled in this study, 39 sites had no visible habitat (classified as ‘Not visible’), due to extremely poor or no visibility at the seafloor. At 10 of these 39 sites, animals were still identifiable in the BRUVs footage. Therefore, a total of 29 sites had neither a visible habitat, nor any associated biodiversity records. These were classified as ‘failed’ deployments and were excluded from further analysis. The total number of successful deployments was therefore 114 sites; these samples were included in the descriptive statistics, species accumulation curves, and diversity indices to reflect the full sampling effort. However, only the 97 samples to which a habitat category could be ascribed were included in the community-level multivariate analyses, and the visualisations of abundance and occurrence relative to habitat and depth, even if no fish species were recorded (MaxN = 0).

For each species, the maximum number of individuals of the same species observed in a single frame, during the 60 min of the sample (referred to as MaxN), was recorded. The MaxN metric eliminates pseudo-replication caused by individuals swimming in and out of the camera’s field of view, and provides a conservative estimate of abundance, especially in high density areas (Cappo et al. 2003; Willis et al. 2000).

### 2.5 Data analyses

A species accumulation curve (SAC) was generated to visualise the completeness of sampling effort (114 samples) using the function ‘specaccum’ and the ‘random’ method to find the mean SAC and its standard deviation from 1000 random permutations of the data (Gotelli & Colwell 2001). The species richness estimators Chao2, Jackknife1, Jackknife2, and Bootstrap were calculated using the function ‘specpool’ for comparison against observed species richness. The diversity indices Species Richness (SR) and Shannon-Weiner diversity (SW) were calculated for 114 samples, and the mean values (± SD) were reported to compare with the estimator range [the lowest value estimator bootstrap (-SE) and the highest value estimator Chao2 (+SE)]. All functions were performed and indices calculated in the R package *vegan* (Oksanen et al. 2026) in R (R Core Team 2024).

The total number of sites where a species was recorded was calculated and reported as N. Frequency of occurrence (FO) was the total number of samples where a species was recorded relative to the total sampling effort (114 samples), reported as a percentage value. For each species: MaxN relative abundance was the sum of all MaxN values divided by the total number of sites sampled (114 sites) and was reported as ‘Mean MaxN All’ to indicate the measure of abundance relative to the total sampling effort. ‘Mean MaxN Present’ was the sum of all MaxN values divided by the number of sites where each species was recorded (present). The standard deviation was calculated to assess the level of dispersion from each mean MaxN value for all species. The distribution of all chondrichthyan species recorded, the most frequently recorded actinopterygian fishes, and two fish species of conservation concern, were plotted using QGIS Version 3.36.3 Maidenhead (qgis.org). The total number of sites where *Haploblepharus pictus* and *Notorynchus cepedianus* were recorded in each habitat type was plotted relative to the total sampling effort within each habitat type, and reported in a bar chart as ‘relative frequency’. These were the only elasmobranchs recorded with a sufficiently high enough numbers of observations to plot relative frequency across habitats.

Variation in MaxN across habitat type was visualised using boxplots showing the median, interquartile range (IQR), and 1.5× IQR whiskers, for five species selected because they were common (klipfishes *Clinus spp.*), abundant (dark shyshark *Haploblepharus pictus*), of regional interest (pelagic goby *Sufflogobius bibartus*), or important linefishery species (Cape gurnard *Chelidonichthys capensis*, Cape seabream *Pachymetopon blochii*). The relationship between MaxN and continuous depth (m) was visualised for each species using a generalised additive model (GAM) smooth (basis dimension k = [4]), fitted using the *mgcv* package (Wood 2011) within ggplot2::geom_smooth(), with 95% confidence intervals. All analyses were conducted in R (version 4.6.1; R Core Team 2024) using the *tidyverse* package (Wickham et al. 2019) for data handling and *ggplot2* (Wickham 2016) for visualisation.

MaxN data from 97 samples that had confirmed habitat descriptions and depth values were square root transformed to promote the influence of rare species and analysed using PRIMER Version 6+ software package (Clarke 1993; Clarke & Warwick 2001). The zero-adjusted Bray-Curtis similarity index (Clarke et al. 2006) was calculated among samples to reflect similarity in relative abundance and species composition.

A one-way Analysis of Similarities (ANOSIM; Clarke 1993) with 999 permutations was performed separately to assess the influence of depth (3 three categories: shallow 1-15 m, medium 16-30 m and deep 31-45 m) and visible habitat (six categories; Appendix I) on species composition and MaxN abundance.

MaxN data were used in a permutational multivariate analysis of variance (PERMANOVA) to test for differences in assemblage structure among habitats and depth categories, with different combinations of these variables and their interactions. PERMANOVA models were evaluated using the pseudo-F statistic with 999 random permutations of the data using the extension software PERMANOVA+ in PRIMER-E v6 (Clarke & Gorley 2006). The similarity percentage (SIMPER) routine identified the contribution of each species towards differences among depth categories and habitat types, and distinguished which species were typically associated with each set of environmental factors (Clarke 1993; Clarke & Warwick 2001).

## 3. RESULTS

A total of 143 samples were collected in this study (Figure 1). The majority of samples were in the shallow (<15 m; 54 sites) and medium (16-30 m; 48 sites) depth categories, with the remaining samples (12) in the deep category (31-45 m). Samples were distributed over unconsolidated sediments, including mud (3 sites), sand (18 sites), and seaweed garden (15 sites); and consolidated sediments, including kelp bed (12 sites), sand-inundated reef (16 sites), and visible reef sites of varying profiles (40 sites); Figure 1 also shows deployment sites where the habitat was not visible (39 sites). Screengrabs from BRUVs footage captured in each visible habitat type are provided in Appendix I. All reporting that follows uses only data from 114 sites (104 with visible habitat and 10 additional samples where habitat was not visible but fauna were) or a subset thereof, collected in August 2022 (23 samples), February 2023 (47 samples), August 2023 (1 sample) and February 2024 (43 samples). Community-level, multivariate analyses used data from 97 sites with visible habitat and accurate depth values.

A total of 17 fish species from 16 families were recorded by the BRUVs (Table 1). Eleven species from 11 families were from the class Actinopterygii (ray-finned fishes, infraclass Teleostei) and five species from four families were from the class Chondrichthyes. One species was an agnathan, or jawless fish, from the class Myxiniformes (sixgill hagfish *Eptatretus hexatrema*). In addition, one marine mammal species (Cape fur seal *Arctocephalus pusillus pusillus*) and two crustacean species (rowing crab *Ovalipes trimaculatus* and West coast rock lobster *Jasus lalandii*) were also recorded, but the analyses that follow focus solely on fishes. A total of 824 individuals were recorded (summed MaxN values), of which 181 individuals were fishes.

**Table 1.**
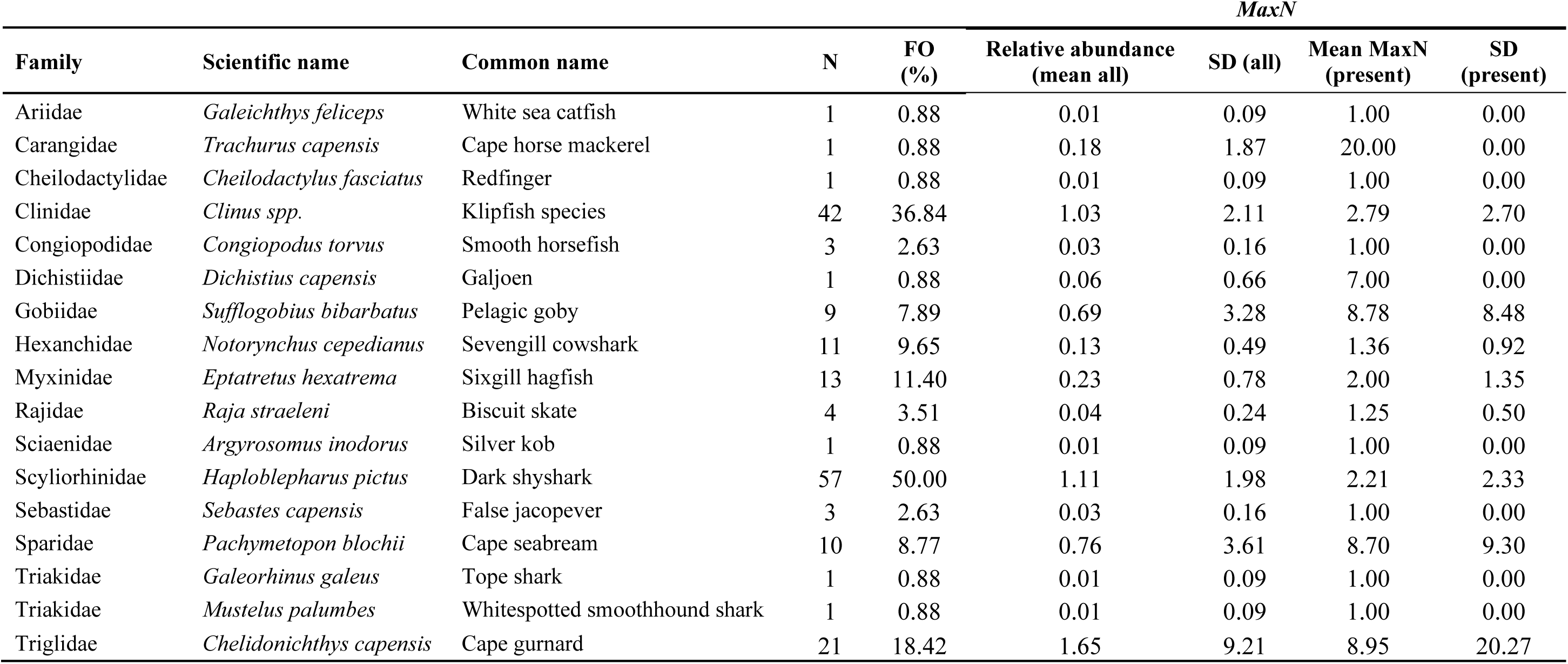
Ichthyofaunal diversity and relative abundance recorded across four sampling fieldtrips conducted between 2022 and 2024 in the Namibian Islands Marine Protected Area (NIMPA). Species are ordered alphabetically by family, and then by species. N denotes the total number of sites where that species was recorded. Frequency of occurrence (FO) is the total number of sites where a species was recorded, divided by the total sampling effort (114 samples) and expressed as a percentage. Relative abundance (mean all) is the sum of all MaxN values for each species relative to the entire sampling effort (divided by 114 samples). Mean MaxN (present) is the sum of all MaxN values for each species relative to the samples where that species was recorded. The standard deviation (SD) is presented for Relative abundance (mean all) and Mean MaxN (present).

### 3.1 Species accumulation curve

The species accumulation curve is shown in Figure 2a. The possible range given by the outputs from four different species richness estimators was as high as 55 species [Chao2 = 37.82 (± 17.20)] and as low as 18.36 [Bootstrap = 19.68 (± 1.31)] (Figure 2b). The non-parametric richness estimator Chao2 (estimate ± SD: 37.82 ± 17.20) represents the upper estimates of species richness to be detected at maximum sampling effort, whereas the richness estimator Bootstrap (estimate ± SD: 19.68 ± 1.31) is closer to the observed total number of species recorded in the survey. The Jackknife 1 (estimate ± SD: 23.94 ± 2.62) and Jackknife 2 (estimate = 30.82) values represent richness estimates between these two extremes (Figure 2b). Mean species richness was 1.58 ± 1.30 (SD) species per sample across 114 samples (range 0-6) and mean Shannon-Weiner diversity was H’ = 0.37 ± 0.42 (SD), where H’ diversity accounts for both species richness and evenness.

**Figure 2:**
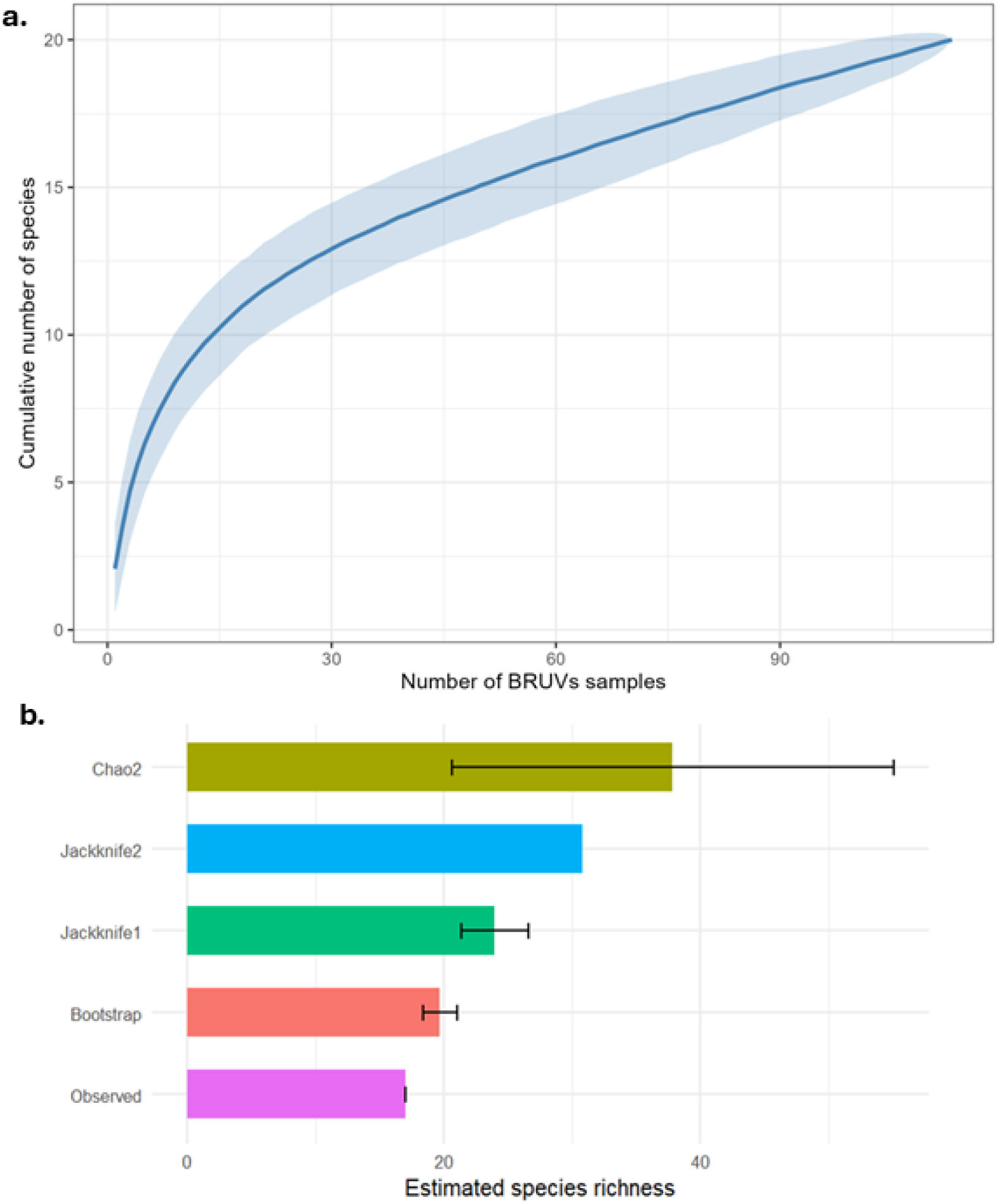
**(a)** Species accumulation curve for fish recorded from 114 samples. The solid blue line shows observed species richness accumulated with sampling effort in this study (method: random permutation, n = 1000 permutations) and ± SD in the shaded blue ribbon. (b) Observed and estimated species richness of fish from 114 samples. Bars show observed species richness (purple), and estimated richness by Chao2, Jackknife 1, Jackknife 2, and Bootstrap calculated using the specpool function in the R package *vegan* (Oksanen et al. 2026). Error bars represent ± 1 SD of each estimator.

### 3.2 Species-specific occurrence, distribution, and MaxN relative abundance

The total records (N) and distribution of all chondrichthyan species recorded (dark shyshark *Haploblepharus pictus*, sevengill cowshark *Notorynchus cepedianus*, tope *Galeorhinus galeus*, whitespotted smoothhound *Mustelus palumbes*, and biscuit skate *Raja straeleni*), the most frequently recorded actinopterygian and agnathan fish species (Cape gurnard *Chelidonichthys capensis*, kilpfishes *Clinus* spp., *Eptatretus hexatrema*, and pelagic goby *Sufflogobius bibarbatus*) and two actinopterigyan fishes of conservation concern, silver kob *Argyrosomus inodorus* and galjoen *Dichistius capensis*, are shown in Figures 3 and 4 relative to the total sampling effort (114 samples).

**Figure 3:**
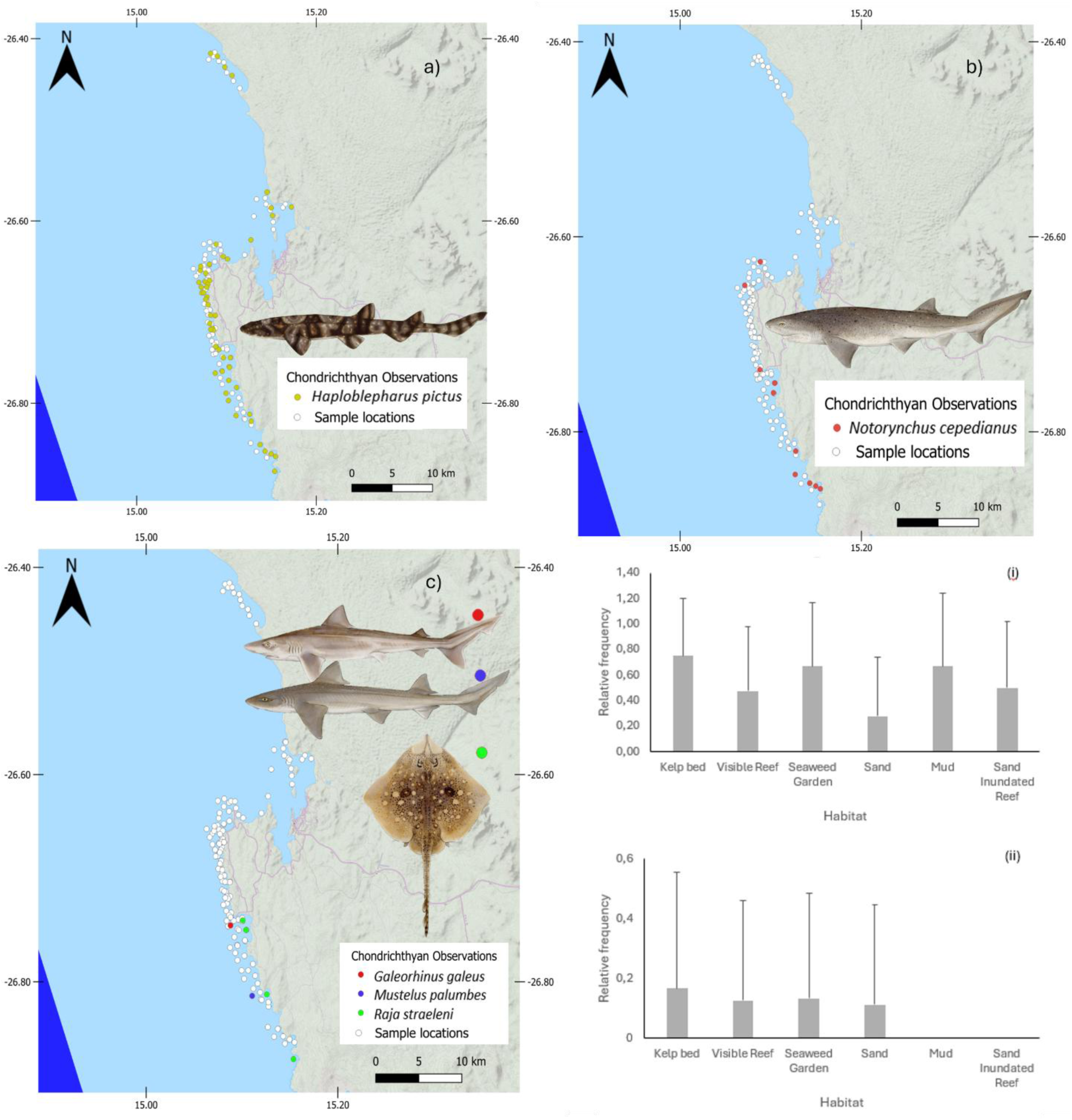
The distribution of the total observations of **a)** Least Concern (LC) dark shyshark (*Haploblepharus pictus*), **b)** Vulnerable (VU) broadnose sevengill cowshark (*Notorynchus cepedianus*) and **c)** Critically Endangered (CR) tope shark (*Galeorhinus galeus*), Least Concern (LC) whitespotted smoothhound shark (*Mustelus palumbes*) and Near Threatened (NT) biscuit skate (*Raja straeleni*) in the Namibian Islands Marine Protected Area (NIMPA) over four sampling fieldtrips conducted between 2022 and 2024. The sites where these species were recorded are shown in coloured circles, and the total sampling effort is shown in white circles. The bar charts show the proportion of observations in each habitat type (+SD) relative to the sampling effort in each habitat type for (i) *H. pictus* and (ii) *N. cepedianus*.

**Figure 4:**
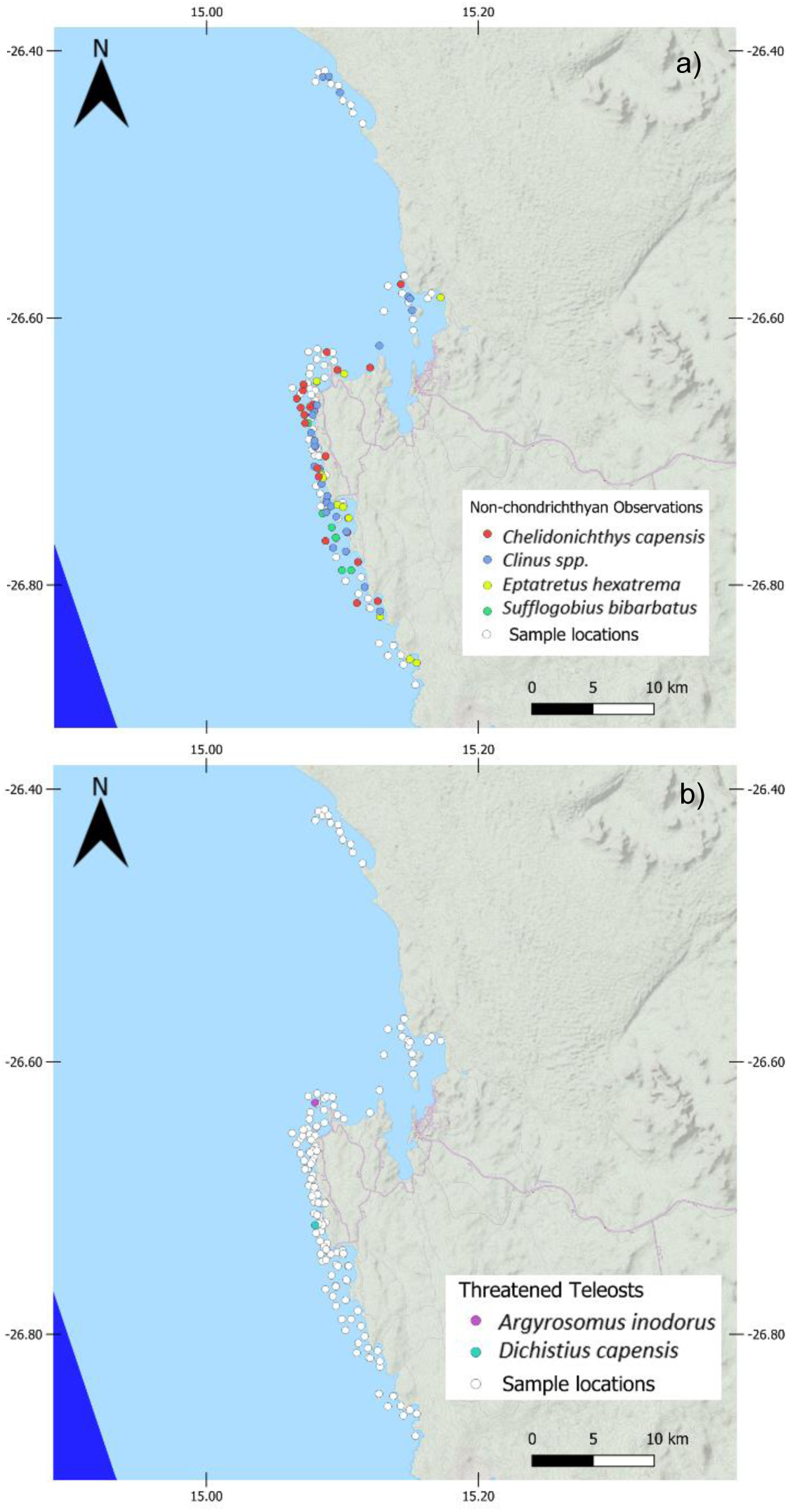
The distribution of the total observations of (a) the four most frequently recorded teleost and agnathan species: Cape gurnard (*Chelidonichthys capensis*), klipfishes (*Clinus spp.*), sixgill hagfish (*Eptatretus hexatrema*) and pelagic goby (*Sufflogobius bibarbatus*) and (b) the two teleost species of conservation concern recorded: silver kob (*Argyrosomus inodorus*), and galjoen (*Dichistius capensis*) in the Namibian Islands Marine Protected Area (NIMPA) over four sampling fieldtrips conducted between 2022 and 2024. The sites where these species were recorded are shown in colour, and the total sampling effort is shown in white circles.

The most frequently recorded species in the survey was the dark shyshark *H. pictus* [FO 50%) (Table 1), which was the most frequently recorded species across all habitat types (75% of all kelp bed sites, 67% of all seaweed sites and 48% of all visible reef sites) and depth categories (63% of shallow, 31% of medium and 67% of all deep samples; Fig. 3). The remaining chondrichthyan species – *N. cepedianus* [FO = 9.65%; Mean MaxN All = 0.13 (± 0.49), *R. straeleni* [FO = 3.51%; Mean MaxN All = 0.03 (± 0.24)], *M. palumbes* [FO = 0.88%; Mean MaxN All = 0.01 (± 0.09)] and *G. galeus* [FO = 0.88%; Mean MaxN All = 0.01 (± 0.09) – were all recorded at lower frequencies (Fig. 3; Table 1).

The most frequently recorded actinopterygian fishes were the klipfishes *Clinus spp.*, sighted at 38% of sites (Table 1; Fig. 4). Most sightings appeared to be *Clinus superciliosus*, with one or two incidences of *Clinus rotundifrons*. These species were recorded at 80% of all seaweed sites, 67% of all kelp bed sites and 35% of all visible reef sites. Relative abundance was highest of all fishes in *Ch. capensis* [Mean MaxN All = 1.65 (± 9.21)], *H. pictus* [Mean MaxN All = 1.11 (± 1.98)] and *Clinus spp*. [Mean MaxN = 1.04 (± 2.11). *Clinus spp.* was frequently recorded in shallow sites, comprising 59% of all shallow samples. *Ch. capensis* was recorded in 19% of all medium samples, and *S. bibarbatus* was recorded in 42% of all deep samples.

### 3.3 Environmental factors and community composition

There was low separation among 97 samples for both factors investigated, depth and habitat type. A one-way ANOSIM showed that depth (Global R = 0.251, p < 0.001) was an important predictor of fish species abundance and community composition, whilst habitat was not significant (Global R = 0.044, p < 0.133). A two-factor PERMANOVA showed that depth was significant (df = 2, Pseudo-F1 = 4.0409, p < 0.001) but habitat (df = 5, Pseudo-F2 = 1.392, p < 0.109) and the interaction between habitat and depth (df = 7, Pseudo-F3 = 1.2445, p < 0.166) were not.

These patterns are visualised for five species (Figure 5). The relative abundance of *H. pictus* was spread across the entire depth range, whilst those of *S. bibartus* and *Ch. capensis* were higher at greater depths. The relative abundances of *Clinus spp*. and *P. blochii* were higher at shallower depths. Likewise, *H. pictus* was recorded in all six habitat types (Figure 6), with the variation in relative abundance highest on visible reef sites. *Clinus spp.* and *P. blochii* were recorded at highest relative abundance in kelp beds and, for the former, on visible reef sites. Mud habitats were where *Ch. capensis* were most abundant, and *S. bibartus* was recorded at a particularly high MaxN value on sand (Figure 6).

**Figure 5.**
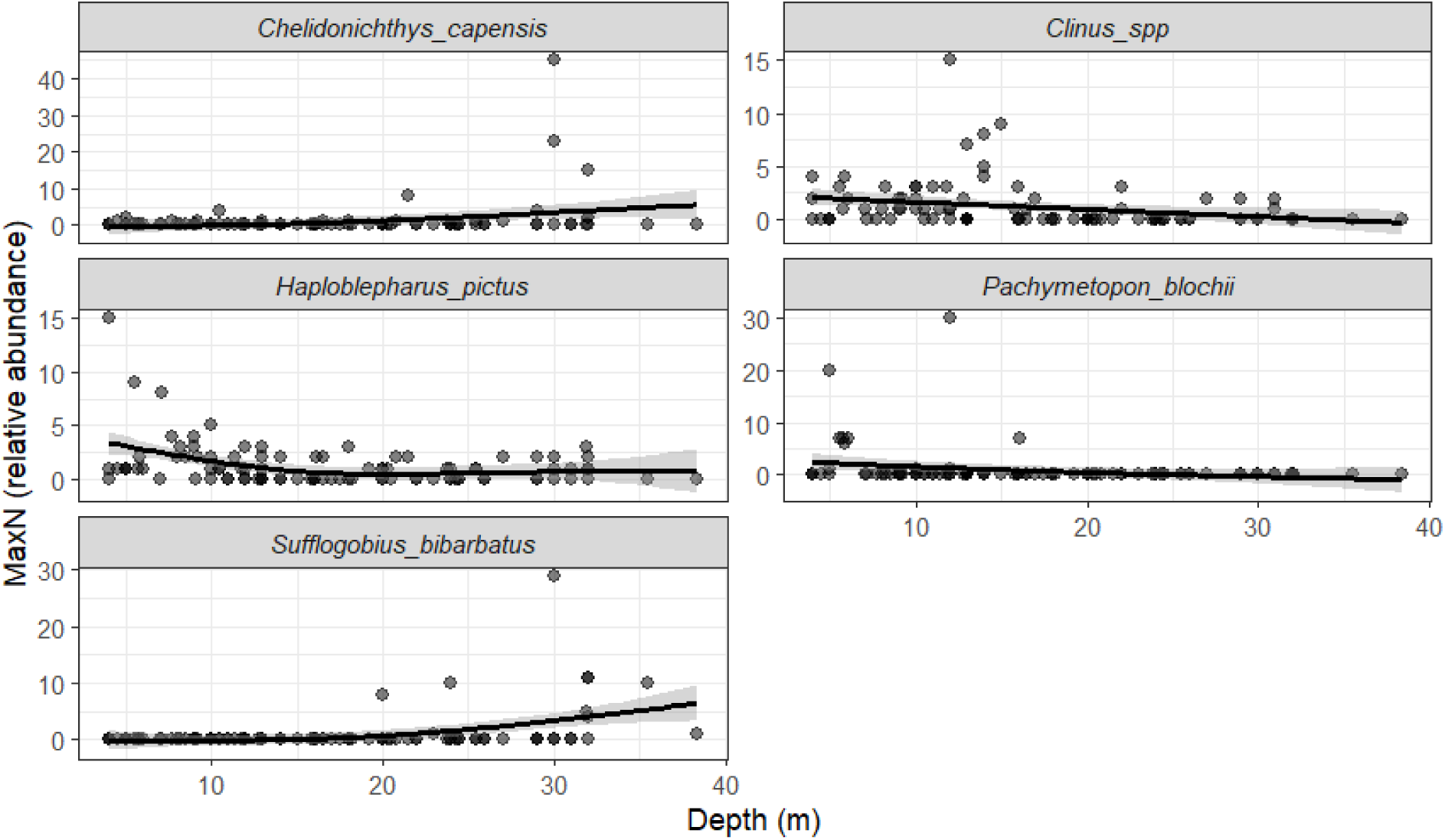
The relationship between relative abundance (MaxN) and depth (m) for Cape gurnard (*Chelidonichthys capensis*), klipfishes (*Clinus spp.*), dark shyshark (*Haploblepharus pictus*), Cape seabream (*Pachymetopon blochii*), and pelagic goby (*Sufflogobius bibarbatus*) recorded from 97 BRUVS deployments. Points show individual sample values; the black line shows a fitted generalised additive model (GAM) smooth (basis dimension k = [4]) with shaded 95% confidence interval. Y-axis scales vary between species panels to accommodate differences in relative abundance.

**Figure 6.**
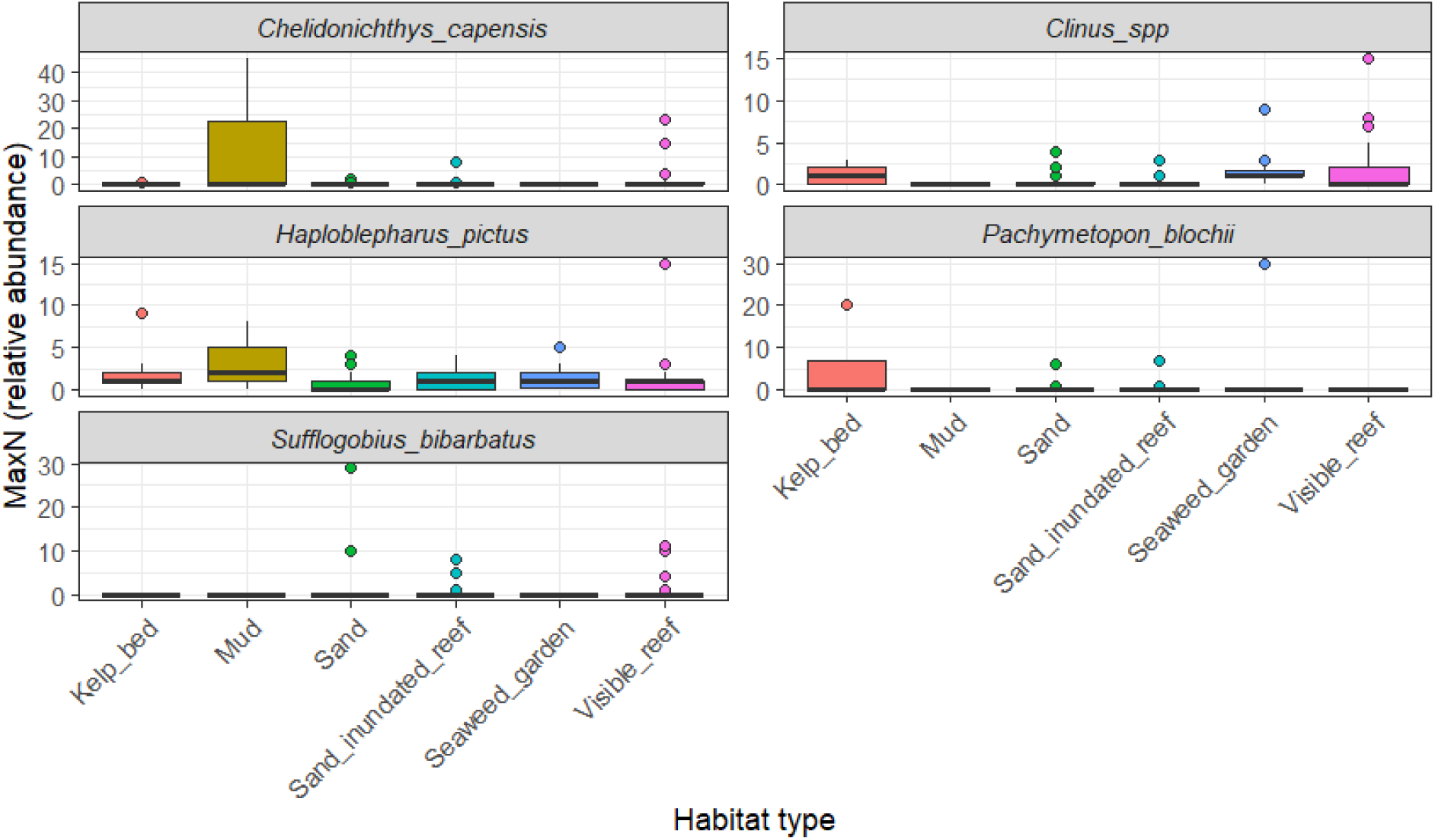
MaxN relative abundance of Cape gurnard (*Chelidonichthys capensis*), klipfishes (*Clinus spp.*), dark shyshark (*Haploblepharus pictus*), Cape seabream (*Pachymetopon blochii*), and pelagic goby (*Sufflogobius bibarbatus*) across six habitat types recorded from 97 samples. Boxes show the median (centre line), interquartile range (box), and 1.5× IQR (whiskers); points beyond whiskers are shown as outliers. Y-axis scales vary between species panels to accommodate differences in relative abundance.

A one-way analysis of similarity percentages (SIMPER) – species contributions – showed that the average similarity was 32.79% between shallow sites, 10.64% between medium depth sites, and 24.19% between deep sites. The SIMPER confirmed that *H. pictus* was ubiquitous and typified all depth categories (40.15% contribution to average similarity in shallow samples, 60.99% in medium depth samples, and 46.46% in deep samples). Additionally, *Clinus spp*. (49.03% contribution to average similarity) were typical of shallow (depths of 0 – 15 m) and medium (16 – 30 m) depth samples (16.20% contribution to average similarity), and *S. bibarbatus* (30.64% contribution to average similarity) was typical of deep samples (31 – 45 m; Fig. 5).

Average similarity was 18.01% between visible reef sites, 36.53% between kelp bed sites, 44.81% between seaweed garden sites, 15.52% between sand-inundated reef sites, 7.02% between sand sites, and 8.61% between mud sites.

The SIMPER confirmed that *H. pictus* was ubiquitous and typified all habitat types (47.46% contribution to average similarity in visible reef sites; 35.26% contribution to average similarity in kelp beds; 37.32% contribution to average similarity in seaweed garden habitat; 68.03% contribution to average similarity in sand-inundated reef sites; 45.13% contribution to average similarity in sand sites; and 100% contribution to average similarity in mud sites).

Additionally, *Clinus spp*. typified visible reef sites (32.77% contribution to average similarity), *E. hexatrema* and *Clinus spp*. typified kelp bed sites (22.40% and 32.12% contribution to average similarity, respectively), and *Clinus spp*. typified seaweed garden sites (58.50% contribution to average similarity) as well as sand sites (33.18% contribution to average similarity).

## 4. DISCUSSION

This represents the first remote subtidal survey of the ichthyofauna of the NIMPA using a single, standardised method to capture baseline biodiversity information. The findings provide insight into ichthyofaunal biodiversity over just two years, combining data from both summer and winter samples. A total of 17 fish species from 16 families were recorded on eight different habitat types, in water depths between 3.4 m and 38.4 m. Three of the species detected are classified as threatened. Depth was an important predictor of abundance of fish species and of community composition, and there were associations, albeit not strong ones, between some species and specific habitat types. The findings suggest that BRUVs can be an effective monitoring approach for the NIMPA and that regular, long-term monitoring using BRUVs would provide insight into the distribution and relative abundance of species of conservation concern, and the effectiveness of the MPA in protecting these habitats and species assemblages.

### 4.1 Species richness and sampling effort

Overall species diversity in this study appears low – 17 species from 16 families – when compared with BRUVs studies in other temperate regions. For example, 27 species belonging to 15 families were recorded in a formerly kelp-dominated coastal ecosystem in Sussex, United Kingdom (Clark et al. 2024); 33 fish species were recorded in a marine reserve and reference location off the eastern coast of Tasmania, Australia, in depths of up to 35 m (Walsh et al. 2017), and 41 teleost species and three elasmobranch species were reported from 280 stereo-BRUVs deployments around the Isles of Scilly, United Kingdom within a similar range of depths to this study (Exeter et al. 2025). Similar studies in South Africa recorded 57 fish species from 30 families in False Bay (De Vos 2021), and 38 species from 14 families (De Vos et al. 2014). Chondrichthyan diversity was also considerably lower in this study - five species from four families - than those reported from studies in South Africa including False Bay (19 species;

De Vos et al. 2015). However, the number of samples collected in False Bay (213) was higher than in this study (143), and the range of depths sampled was also far greater: 84 m in False Bay and only 38.4 m in this study. Whilst both surveys were in what are considered temperate waters, False Bay is located at the confluence of the Benguela and the Agulhas Currents (Day 1970), with the overlap of two major biogeographic regions giving rise to higher overall diversity incorporating species typical of the cooler west coast alongside those typical of the warmer south coast (Griffiths et al. 2010). In addition, the Lüderitz upwelling cell, driven by strong, perennial southerly winds, acts as an environmental barrier to the movements of some species and the transport of plankton and fish larvae (Duncombe Rae 2005; Lett et al. 2007).

There are no published BRUVs data from South Africa’s west coast or from the Angolan coast and thus the diversity and relative abundance indices described here could not be compared with other sites within the Benguela Current region. Several species known to be captured from shore by recreational anglers in Lüderitz or elsewhere along the Namibian coast were not recorded on the BRUVs. Silver kob *Argyrosomus inodorus* and West coast steenbras *Lithognathus aureti* are the primary recreational and commercial linefish species targeted in the northern Benguela (Holtzhausen et al. 2001), but just one detection of one of these species was recorded during the study. Chondrichthyan species known to be present in the area include the spotted gully shark *Triakis megalopterus*, bronze whaler shark *Carcharhinus brachyurus* and bluntnose guitarfish *Acroteriobatus blochii* (Leeney 2024b; Leeney et al. 2024; Rogers et al. 2022). Likewise, large numbers of white skate *Rostroraja alba* egg cases have been observed on several beaches around Lüderitz (RHL pers. obs.), but this species was not present in the BRUVs samples. Although indicative only of the larger fish species tagged by recreational anglers, data from the Oceanographic Research Institute (South Africa) Cooperative Fish Tagging Project, which includes records from Namibia over three decades (up to c. 2013), indicate the presence of at least 17 chondrichthyan species and 20 teleost species (RH Leeney unpubl. data). The diversity recorded in this study is thus a fraction of the overall biodiversity in Namibia’s coastal waters. Most southern African BRUVs surveys have concentrated on reef habitats further to the east, in the more biodiverse warm-temperate waters which do not provide a valid point of comparison given the greater biodiversity of that region (Bernard 2012; Bernard & Götz 2012; Sanguinetti 2013; De Vos et al. 2014; Heyns-Veale 2016; Martinez et al. 2024; Parker et al. 2016). In contrast, there remains a paucity of baseline information on the ichthyofauna on South Africa’s cool-temperate west coast (Gazeau et al. 2026).

The lack of baseline data for Namibian subtidal habitats and marine assemblages makes it challenging to assess whether the low diversity described in this study indicates a decline in elasmobranchs and in reef and predatory fish species that might be considered for inshore waters of southern Africa. The species accumulation curve and richness estimates reported here suggest that 114 BRUVs deployments undersampled the fish community, with the Chao2 estimator suggesting almost double the number of potential species should be detected. This is especially likely if many species are not abundant and are spatially dispersed, and more so if they preferentially inhabit waters further from the zone most impacted by humans (Lüderitz Bay). However, the Chao2 estimator accounts for incidents of rare species in the dataset (Chao et al. 2009; Chao & Chiu 2016) and therefore represents the upper limit of what effort is actually required; in testing the methods, Chao et al. (2009) reported that while estimates could range up to 10.67 times the original sample, less effort is needed to detect 90% of the species.

### 4.2 Relative abundance

The most abundant and ubiquitous chondrichthyan in this study was *Haploblepharus pictus*, which is the west coast counterpart to the puffadder shyshark *H. edwardsii*, the most prevalent species in records from temperate South African BRUVs surveys (De Vos et al. 2015). The most prevalent large shark was *Notorynchus cepedianus*, which was recorded at 11 sites. This species has a very diverse diet including other chondrichthyans such as *Haploblepharus* spp. and *Raja* spp.; teleosts, delphinids, Cape fur seals and cephalopods (Ebert 1991). This broad diet may allow *N. cepedianus* to persist in areas where other chondrichthyans with more specific diets cannot.

Whilst species diversity is not expected to be comparable between southern Namibia and False Bay, South Africa, relative abundance might be expected to be so, particularly in an upwelling regime. In this case, apart from *N. cepedianus*, larger-bodied sharks and other large predatory fishes were largely absent or in low abundance in the NIMPA (only one record each for two medium-size shark species, *Galeorhinus galeus* and *Mustelus palumbes*). Where sharks have been subject to intense fishing pressure and habitat loss in the Arabian Gulf, relative abundance was reportedly low despite a large survey area and extensive sampling effort (Jabado et al. 2018). Fishing pressure in the region prior to its designation as an MPA, combined with overfishing of key prey species such as *Sardinops sagax* (Erasmus et al. 2021), and continued degradation of coastal habitats and possible changes to ecosystem structure caused by diamond mining (Pulfrich et al. 2003; Pulfrich and Branch 2014), may have similarly reduced predator populations to reflect the low abundance reported here. However, not all species were reported in low relative abundance. *Chelidonichthys capensis*, *Pachymetopon blochii*, *Sufflogobius bibartus*, *Clinus* spp. and even *N. cepedianus* were all recorded at relative abundances higher than reported in studies elsewhere. Indeed, *N. cepedianus* (relative abundance of 0.13 in this study) was recorded at 0.05 in False Bay, South Africa (De Vos et al. 2015) where they have been known to aggregate (Engelbrecht et al. 2020). It is also important to consider the type of species that might have highest expected biomass on the Namibian coast; shoaling, pelagic species are not well sampled by benthic BRUVs and this method may therefore underrepresent the ichthyofaunal community that is most prevalent in the region.

### 4.3 Environmental predictors of species composition

Although the geographic scope of the study and range of depths sampled was limited, a wide range of habitat types were documented. Habitat type, depth and season have all been found to be key predictors of species composition (e.g. De Vos et al. 2015; Walsh et al. 2017). It was not possible to assess differences between seasons due to the small dataset collected during winter, but depth was an important predictor of the abundance of fish species and of community composition. *H. pictus* was ubiquitous and typified all depths and habitat types, whilst several species were characteristic of specific habitats or depth categories. The most frequently recorded species – *H. pictus* and *Clinus spp.* - were found on most habitat types (but most frequently in kelp bed sites), and at all depths (but most frequently in the shallow depth category). This suggests that these species are generalists and can adapt to a range of habitats and conditions. *S. bibarbatus* was typical of deep samples, although the total number of deep samples in this study was low. Given that this species is now a key component of the marine food web in southern Namibia (Utne-Palm et al. 2010) and an important prey species for several threatened species that inhabit the NIMPA including African penguins *Spheniscus demersus* (Critically Endangered; Birdlife International 2024; Ludynia et al. 2010), the identification and consistent monitoring of habitats for this species should be an important component of long-term monitoring. The lack of samples from even deeper areas may have excluded species that may be specific to those habitats, and future studies in the NIMPA should aim to collect BRUVs samples from a greater range of depths.

Biodiversity on sand tends to be more complex to measure and monitor, as species diversity is less evenly distributed and sand-associated species have variable responses to MPA protection at different scales (Fetterplace 2017). This often means that, in order to capture diversity associated with sand habitats (and other soft sediments), wider survey coverage is required than for reef habitat (De Vos 2021). This study, however, points to the importance of ensuring that sand and other soft sediments are included in biodiversity monitoring and protection objectives for the NIMPA: *G. galeus* (Critically Endangered) was only recorded on sand, and several other species were reported from sandy habitats, including *R. straeleni*, *N. cepedianus* and *H. pictus*. This result reflects similar findings for the same species in both temperate and subtropical South Africa (De Vos 2021; Ferreira et al. 2023) and emphasises the importance of soft sediments to many chondrichthyans, especially skates, rays and chimaeras, and to some threatened teleost fishes (Martin et al. 2012; Martins et al. 2020).

Habitat classes are increasingly being used as the basis for designing MPAs to ensure adequate representation of the total biodiversity of an area (Ward et al. 1999, Mumby & Hastings 2008, Rees et al. 2018). A study in Jervis Bay, Australia, used BRUVs to assess spatial variability in demersal and mid-water fishes at multiple scales, to determine whether habitat classes are appropriate surrogates for temperate fishes (Rees et al. 2018). The findings revealed that there was a distinct assemblage of demersal fishes associated with each habitat class, driven by strong habitat associations for many families and species, whilst the mid-water fish assemblage and certain demersal families, such as habitat generalists (e.g. some sparids), showed no selectivity for specific habitat classes. Further data collection in the NIMPA may reveal whether there are particular habitats that species of conservation and management concern associate with. Determining the best scale at which to delineate habitat types, based on the community in focus, may also yield different results. For example, differences in invertebrate epifauna may be more strongly distinguished at finer habitat classification scales than demersal fish, and mobile predators like sharks (De Vos 2021). Mapping such habitats may then provide a useful first step in prioritising areas for monitoring in and future zonation of the NIMPA.

### 4.4 Threatened species

Three of the species recorded during the study are considered threatened by the IUCN Red List. *G. galeus* was recorded only once; this species has undergone declines throughout its range as a result of targeted and accidental captures in small-scale and industrial fisheries (Walker et al. 2020). More research, potentially involving acoustic telemetry, is urgently needed to better understand the distribution of and threats to *G. galeus* in Namibian waters, and to assess whether the NIMPA provides important habitat for the species. *N. cepedianus* (Vulnerable; Finucci et al. 2020) appeared to be relatively common within the sampled area and the NIMPA may therefore provide important habitat for this species. *A. inodorus* (Vulnerable; Fennessy & Winker 2020) was only recorded once during this study, but this species is typical of the surf zone where it occurs west of Cape Agulhas in South Africa, and juveniles favour sandy bays shallower than 50 m (Heemstra & Heemstra 2004). This study’s sampling distribution may have excluded some of the preferred habitat for this species. Line fishing is the major threat for this species, which was historically overfished in Namibian waters (Kirchner 1998; Fennessy & Winker 2020). The presence of *Dichistius capensis* (not yet been assessed for the Red List) is of note, as it occurs in two disjunct stocks with some exchange between them: a western stock along the coast of Namibia and the west coast of South Africa, and a southern stock on the southern coast of South Africa (Attwood & Bennett 1994, Attwood & Cowley 2005). The Namibian stock has not been assessed, but the South African stock is considered to have collapsed, and the population is estimated to be at less than 20% of pristine levels (Mann 2013).

### 4.5 Threats

The food web in the NIMPA and throughout Namibian waters is fundamentally different today from its state in the 1960s, as is the case for many of the historically highly productive marine ecosystems such as those in northwest Africa, Peru, Chile and California. Historically, sardine *S. sagax* was an important forage species for many pelagic species in the northern Benguela Upwelling System, but the sardine population collapsed in the 1970s and 1980s due to a combination of overfishing and ecosystem change and variability in the region (Crawford et al. 1987; Boyer et al. 2001). This collapse of a key prey species changed the structure and function of the ecosystem, which became dominated by jellyfish, pelagic goby, hakes (*Merluccius* spp.) and Cape horse mackerel *Trachurus capensis* (van der Lingen et al. 2006) and may have had far-reaching impacts on the entire northern Benguela Upwelling ecosystem through feeding linkages (Erasmus et al. 2021). Other environmental factors such as Benguela Niños, anomalous warming events associated with poleward intrusions of warm, saline tropical surface waters along the Namibian and Angolan coastlines (Shannon et al. 1986), and climate change-induced warmer temperatures in northern Namibian waters alongside some temperature decreases in the south (Lima & Wethey 2012), may also have caused or contributed to changes in the availability of prey species. Some predators were able to switch diets from sardines to pelagic gobies after the collapse of the sardine populations (Utne-Palm et al. 2010; Erasmus et al. 2021), but others may not have had this flexibility. The paucity of information on the diets of many predator species and a lack of historical data on local marine biodiversity, prior to the collapse of the sardine population, make it challenging to assess whether the ichthyofauna of the NIMPA has been significantly altered by this change in the food web, but the presence of primarily smaller fish species, the mesopredator *H. pictus* and the scavenger *Eptatretus hexatrema*, and the absence or rarity of many larger predatory fishes known to inhabit Namibian coastal waters, suggests a prevalence of lower trophic levels. The rarity of large predatory fishes cannot on its own be ascribed to population declines; it may simply be a result of the sampling coverage or some bias in the method. However, very few of the natural predators of *H. pictus* (either chondrichthyan or teleost) were recorded in this survey and where they were (e.g. *N. cepedianus*), they were at low abundance relative to the prey species. This may suggest that there have already been declines in large predators in the region, causing mesopredator release, but it is also possible that the abundance of these smaller sharks has always been high (relative to that of other shark species) in southern Namibia, and that this simply has never previously been documented.

The NIMPA has to date been an MPA without an active management plan and with no consistent monitoring of species other than those of commercial interest. Despite its designation as an MPA, it is subject to ongoing diamond mining along the coast and in the inshore marine environment, which involves cutting kelp, sucking up gravel from the seafloor, which is sorted on the shore and then deposited intertidally, and uncovering and overturning subtidal boulders (Pulfrich et al. 2003). This mining alters the biological and physical characteristics of the habitat, impact the seabed and thus the benthic fauna (Rogers and Li 2002), has visible impacts on water visibility and sediment load (RH Leeney pers. obs.) and likely impacts the biodiversity in and adjacent to mined areas. At least two salmon aquaculture developments have been approved for development inside the NIMPA^1^, and such activities have been documented to pose a range of threats to the marine environment including the escape of non-native species, localised changes in the physico-chemical properties of benthic sediments and loss of macrofaunal biodiversity (Keeley et al. 2015; Buschmann et al. 2006).

While it is often assumed that recreational fishing has negligible impact on fish populations in comparison to commercial fishing, research has shown that this is not always the case. Recreational angling can slow the recovery of some species that have previously been impacted by commercial fishing (e.g. Dainys et al. 2022). In South Africa, angling has been shown to have a significant impact on some fish populations and even catch-and-release fishing can have an impact, with increased post-release mortality evident for more sensitive fish species (Mann et al. 2018). This has led to the recommendation that catch-and-release shore-based angling by members of the public is not compatible with MPAs zoned as no-take areas.

### 4.6 Recommendations

This study demonstrated that this method works in the Namibian context and has collected novel data that provide an insight into the marine biodiversity of the NIMPA. However, the findings are not representative of the entire NIMPA due to the study’s limited spatial scope and small sample size. Along the NIMPA’s westernmost boundary, there are reportedly water depths of up to 165 m; whilst the northernmost and southernmost parts of the NIMPA are less disturbed than the area around Lüderitz, which undergoes dredging for the port, and around which there is frequent vessel traffic including small recreational vessels, commercial fishing vessels and cruise ships (RH Leeney pers. obs.). The marine habitats beyond the area sampled in this study are poorly described and may harbour many additional species not documented hereby this study. Extending monitoring to a greater range of depths and into areas further to the north and south, and using pelagic BRUVs to complement the data from benthic systems, are priorities for future work. Future efforts to sample in the NIMPA in both summer and winter, and to collect temperature data for each BRUVs deployment, would provide insight into changes in community structure and the relative abundance of species between seasons, and the role temperature may play in determining species distributions. Extracting length measurements from stereo-BRUVs data will also provide valuable insights into size classes and population structuring of key species.

Using BRUVs alone will not comprehensively document the NIMPA’s biodiversity or provide all the necessary data for management. Typically, benthic BRUVs do not adequately sample pelagic and shoaling species (like mackerel and sardines). As such, these records are opportunistic and the relative abundance measures of these species would be better monitored using fisheries surveys or potentially, pelagic BRUVs. Several fish species documented in this study are important fisheries targets or bycatch, in parts of their range: the mackerel species, gurnard, seabream and kob. For instance, *P. blochii* is an important component of the commercial line fishery in South Africa (Mann 2013); this species was recorded in 9% of samples in this survey, and its relative abundance was fourth highest in the survey. The long-lived, fast-growing *C. capensis* is considered a r-selected generalist species (McPhail et al. 2001) that is caught as bycatch in both inshore and offshore demersal trawl fisheries in Southern Africa (Smale and Badenhorst 1991, Japp et al. 1994); it was recorded in 18% of samples and had the highest relative abundance of all teleost fishes recorded. The ability to monitor species of commercial importance that may have experienced declines as a result of fishing pressure, or may continue to decline in areas where they are unprotected, allows for the efficacy of the protection provided by an MPA to be assessed as part of a suite of fisheries management tools.

Effective conservation of marine vertebrates, especially highly mobile species, relies on a good understanding of their patterns of habitat use, movements and migrations (e.g. Daly et al. 2023; Kraft et al. 2023a; Doherty et al. 2017). Some large marine vertebrate species can be highly mobile and can undertake long migrations (e.g. Rogers et al. 2022), and movement data can provide insight into the habitats a species uses, as well as the threats it may encounter as it moves from one region to another, both of which have direct implications for conservation (Lennox et al. 2023; Dwyer et al. 2020). Understanding the movement behaviour of species of conservation interest is thus essential to improve the protection offered by established MPAs and to more effectively design new MPAs (Dwyer et al. 2020). Conventional tag-and-release programmes such as the ORI-CFTP (Dunlop et al. 2013) or acoustic tracking, which has been successfully piloted at a small scale within the NIMPA (Leeney et al. 2024), would provide valuable insight into the movements and home ranges of species of conservation concern, informing fisheries management and marine spatial planning (Lennox et al. 2023). The teleosts of conservation concern highlighted by this study (*A. inodorus* and *D. capensis*) are often typical of shallow, turbid water and as such, are less likely to be captured by BRUVs sampling. Indeed, water visibility is perhaps the greatest limitation to the use of BRUVs (Jabado et al. 2018), and poor visibility in many of this study’s samples may have reduced the likelihood of detecting some species. An alternative method for sampling these habitats (e.g. seine net sampling of nearshore and surf zone communities; eDNA) should also be incorporated into the NIMPA’s monitoring plan. However, BRUVs help to capture community composition, habitat and ecosystem-level changes over time, and thus they provide valuable insight into how ecosystem structure and function may be changing. Should strategies for managing and restoring the marine fauna within the NIMPA be implemented, monitoring data collected by stereo-BRUVs surveys could also provide measurements of species-specific recovery patterns and track the efficacy of such strategies (Jabado et al. 2018).

### 4.7 Conclusions

This study has demonstrated the value of BRUVs for monitoring marine biodiversity in Namibian waters. The data have provided a baseline species list, which allowed for species of conservation concern and potential focal species for longer-term monitoring efforts to be highlighted and added to the NIMPA management plan (currently in review with the Namibian Ministry of Agriculture, Fisheries, Water and Land Reform). The results indicate that in comparison to findings from other studies in temperate coastal waters, the NIMPA appears to have low levels of ichthyofaunal biodiversity and abundance, although the underlying reasons for this result are unknown. The lack of historical data for the region prevents a complete understanding of the status and health of the areas sampled. Developing an active management approach and potentially directed restoration activities within the NIMPA, rather than simply protecting the area in its current state may, over time, reveal whether the area can support a greater diversity of marine fauna. The results presented here provide a baseline against which the results of future monitoring can be compared, but monitoring should be extended to other parts of the NIMPA for a more complete picture.

Previous research has suggested that, because of vulnerability of pelagic ecosystems to human pressures such as industrial fisheries, the protection and recovery of large predatory, pelagic fishes require the creation of highly protected areas in remote locations (Letessier et al. 2024). The offshore waters of the NIMPA, as well as its coastal waters alongside the inaccessible stretches of coastline, may provide such remote conditions. Although these are by their nature more difficult to survey, they may provide valuable refuges for threatened species, and it will be essential to monitor and protect biodiversity in such regions in the future.

## ACKNOWLEDGEMENTS

All BRUVs videos collected as part of this study have been provided to the Namibian Ministry of Agriculture, Fisheries, Water and Land Reform. A copy of the dataset has been stored on servers in the South African Institute for Aquatic Biodiversity (SAIAB), Makhanda, South Africa. We are grateful to SAIAB/ ACEP for providing the BRUVs equipment.

This work was conducted under a research permit from the National Commission on Research, Science and Technology; Certificate number RCIV00012021, authorisation number AN202202001. Ethical clearance for this work was granted by the University of Namibia, Ethical Clearance Reference Number SNC0001.

Many thanks to our skippers, Stefan and Heiko Metzger; to Finlay Bell, Kalimukwa Manyando, Ndamononghenda Mateus, Priskilla Nghaangulwa and Arariky Shikongo for assistance with fieldwork, and to the land-based safety support team: R. Braby, F. Chase, S. Kahunda, S. Matjila, and J. Nowotes. We are also grateful to Dr Boyd Escott for assistance with cartography, and to James Seager (SeaGIS) and Sarah Brien and Marti J. Anderson (PRIMER-e (Quest Research Ltd)).

This work was funded by grant number G22-212-6212 from the Shark Conservation Fund, a project of Rockefeller Philanthropy Advisors.

Illustrations by Alexis Aronson.

**Appendix 1:**
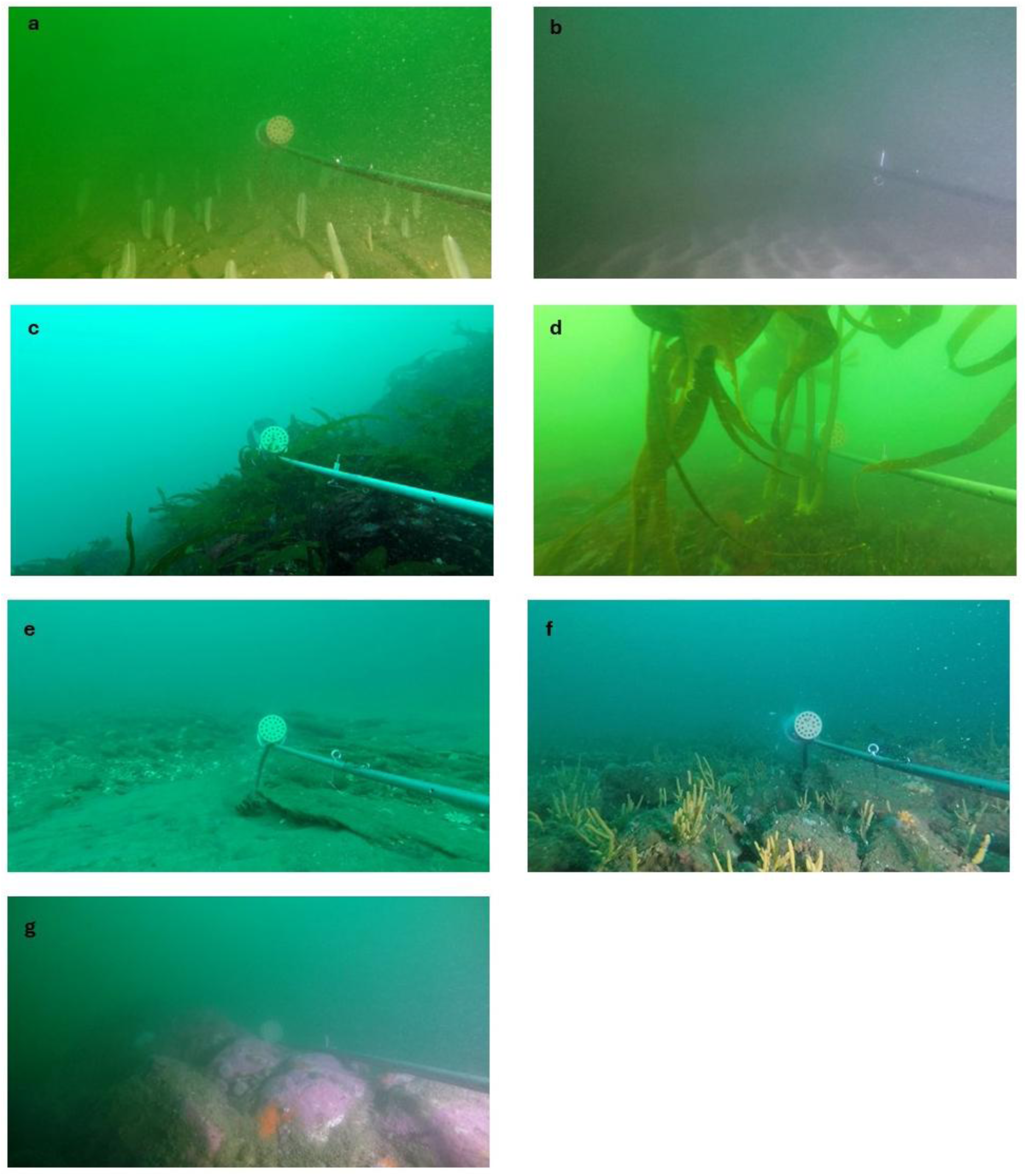
Screen grabs from BRUVs footage collected in the NIMPA, showing features of the predefined habitat types. (a) Mud, (b) sand, (c) seaweed garden, (d) kelp bed, (e) sand-inundated reef, and the visible reef category which includes (f) low profile rocky reef and (g) high profile rocky reef.

## Footnotes

1 Jiménez (2024) https://weareaquaculture.com/news/aquaculture/norwegian-company-secures-license-for-offshore-salmon-farming-in-namibia

